## Supplementary material for "Biochemical principles of miRNA targeting in flies": Table S1

**Table S1. RNA and DNA oligonucleotides used in this study**

| **Ago1 loading** | **Sequence**<br>Seed; p indicates 5′ monophosphate |
|---|---|
| *let7* guide strand | pUGA GGU AGU AGG UUG UAU AGU |
| *let7* passenger strand | UAU ACA AUG UGC UAG CUU UCU |
| *bantam* guide strand | pUGA GAU CAU UUU GAA AGC UGA UU |
| *bantam* passenger strand | CCG GUU UUC GAU UUG GUU UGA CU |
| miR-184 guide strand | pUGG ACG GAG AAC UGA UAA GGG C |
| miR-184 passenger strand | CCU UAU CAU UCU CUC GCC CCG |
| miR-11 guide strand | pCAU CAC AGU CUG AGU UCU UGC |
| miR-11 passenger strand | CAA GAA CUU UCU CUG UGA CCC G |
| **Ago1•miRNA purification** | **Sequence**<br>RNA, DNA; m, 2′-*O*-methyl ribose; p, 5′ phosphate;<br>Bio, Biotin-6-carbon spacer |
| Capture Oligo to affinity purify Ago1•*let-7* | Bio-mAmUmA mGmAmC mUmGmC mGmAmC mAmAmU mAmGmC mCmUmA mCmCmU mCmCmG mAmAmC mG |
| DNA competitor to elute Ago1•*let-7* | Bio-CGT TCG GAG GTA GGC TAT TGT CGC AGT CTA T |
| Capture Oligo to affinity purify Ago1•*bantam* | Bio-mAmUmA mGmAmC mAmCmU mUmGmU mUmUmC mCmCmU mUmUmG mAmUmC mUmCmC mGmCmC mCmG |
| DNA competitor to elute Ago1•*bantam* | Bio-CGG GCG GAG ATC AAA GGG AAA CAA GTG TCT AT |
| Capture Oligo to affinity purify Ago1•miR-184 | Bio-mAmUmA mGmUmA mAmAmC mCmAmU mCmAmU mCmCmA mUmCmC mGmUmC mCmCmG mAmAmC mG |
| DNA competitor to elute Ago1•miR-184 | Bio-CGT TCG GGA CGG ATG GAT GAT GGT TTA CTA T |
| Capture Oligo to affinity purify Ago1•miR-11 | Bio-mAmAmA mAmAmA mCmCmU mAmAmC mUmAmU mAmCmC mUmGmU mGmAmU mAmAmA mAmAmG |
| DNA competitor to elute Ago1•miR-11 | Bio-CTT TTT ATC ACA GGT ATA GTT AGG TTT TTT |
| **Ago1•miRNA quantification** | **Sequence**<br>p, 5′ phosphate |
| RNA probe to quantify total concentration of Ago1•*let-7* by Northern Blot | pGAU ACU AUA CAA CCU ACU ACC UCA ACC U |

| | |
|---|---|
| RNA probe to quantify total concentration of Ago1•*bantam* by Northern Blot | pGUU AAU CAG CUU UCA AAA UGA UCU CAU AGA |
| RNA probe to quantify total concentration of Ago1•miR-184 by Northern Blot | pGUA GCC CUU AUC AGU UCU CCG UCC AAU UA |
| RNA probe to quantify total concentration of Ago1•miR-11 by Northern Blot | pGUA GCA AGA ACU CAG ACU GUG AUG AAG U |
| RNA target to quantify active concentration of Ago1•*let-7* by double-filter binding | pGAA AAA AAA AAA AAA UCU ACC UCU AAA U |
| RNA target to quantify active concentration of Ago1•*bantam* by double-filter binding | pAAA AAA AAA AAA AUU UUU GAU CUC UAA AU |
| RNA target to quantify active concentration of Ago1•miR-184 by double-filter binding | pGAA AAA AAA AAA AAA AAU CCG UCC UAA AU |
| RNA target to quantify active concentration of Ago1•miR-11 by double-filter binding | pGAA AAU UUU AAA AAA UCU GUG AUA AAA U |
| **Substrates for competition assays** | **Sequence**<br>m, 2′-*O*-methyl ribose; ps; phosphorothioate; complementary to guide |
| Target to *let-7* with g2–21 complementarity | pGAU ACU AUA CAA CmCpsmU ACU ACC UCA ACC U |
| Target to *let-7* with g2–21 complementarity and a G:U pair at position 4 | pGAU ACU AUA CAA CmCpsmU ACU ACU UCA ACC U |
| Target to *let-7* with g2–21 complementarity and G:U pairs at positions 4–5 | pGAU ACU AUA CAA CmCpsmU ACU AUU UCA ACC U |
| Target to *let-7* with g2–8 complementarity and mismatches at positions 4–5 | pGAA AAA AAA AAA AmApsmA UCU AAA UCA AAA U |
| Target to *let-7* with g2–8 and g13–16 complementarity and mismatches at positions 4–5 | pGAA AAA AAA CAA AmApsmA UCU AAA UCA AAA U |
| Target to *let-7* with g2–8 and g12–17 complementarity and mismatches at positions 4–5 | pGAA AAA UUA CAA CmApsmA UCU AAA UCA AAA U |

| | |
|---|---|
| Target to *let-7* with g2–21 complementarity and mismatches at positions 4–5 | pGAU ACU AUA CAA CmCpsmU ACU AAA UCA ACC U |
| **Substrates for in vitro cleavage** | **Sequence**<br><u>Seed</u>; m, 2′-*O*-methyl ribose; ps; phosphorothioate;<br>complementary to guide |
| Target to *let-7* with g2–20 complementarity | GAG UUC UAC AGU CCG ACG AUC CUA UAC AAC CUA <u>CUA CCU C</u>AU GGA AUU CUC GGG UGC CAA |
| Target to *let-7* with g2–8 complementarity | GAG UUC UAC AGU CCG ACG AUC AAA AAA AAA AAU <u>CUA CCU C</u>AU GGA AUU CUC GGG UGC CAA |
| Target to *let-7* with g4–14 complementarity | GAG UUC UAC AGU CCG ACG AUC AAA AAA AAC CUA CUA CCA AUU GGA AUU CUC GGG UGC CAA |
| Target to *let-7* with g5–15 complementarity | GAG UUC UAC AGU CCG ACG AUC AAA AUC AAC CUA CUA CAA AUU GGA AUU CUC GGG UGC CAA |
| Target to *let-7* with g4–14 complementarity and modified t10,t11 | GAG UUC UAC AGU CCG ACG AUC AAA AAA AAC mCpsmUA CUA CCA AUU GGA AUU CUC GGG UGC CAA |
| Target to *let-7* with g5–15 complementarity and modified t10,t11 | GAG UUC UAC AGU CCG ACG AUC AAA AUC AAC mCpsmUA CUA CAA AUU GGA AUU CUC GGG UGC CAA |
| **RBNS** | **Sequence**<br>RNA, DNA |
| RBNS RNA input pool | pGAG UUC UAC AGU CCG ACG AUC NNN NNN NNN NNN NNN NNN NNU GGA AUU CUC GGG UGC CAA |
| 5′-end blocking cDNA oligonucleotide #1 | GTC GGA CTG TAG AAC TC |
| 3′-end blocking cDNA oligonucleotide #1 | TTG GCA CCC GAG AAT |
| 5′-end blocking cDNA oligonucleotide #2 | CGG ACT GTA GAA CTC |
| 3′-end blocking cDNA oligonucleotide #2 | TTG GCA CCC GAG A |
| RT primer | CCT TGG CAC CCG AGA ATT CCA |
| PCR Forward primer | AAT GAT ACG GCG ACC ACC GAG ATC TAC ACG TTC AGA GTT CTA CAG TCC GA |
| Multiplexing PCR Reverse Primer PCRId1 | CAA GCA GAA GAC GGC ATA CGA GAT CGT GAT GTG ACT GGA GTT CCT TGG CAC CCG AGA ATT CCA |
| Multiplexing PCR Reverse Primer PCRId2 | CAA GCA GAA GAC GGC ATA CGA GAT ACA TCG GTG ACT GGA GTT CCT TGG CAC CCG AGA ATT CCA |
| Multiplexing PCR Reverse Primer PCRId3 | CAA GCA GAA GAC GGC ATA CGA GAT GCC TAA GTG ACT GGA GTT CCT TGG CAC CCG AGA ATT CCA |

| | |
|---|---|
| Multiplexing PCR Reverse Primer PCRId4 | CAA GCA GAA GAC GGC ATA CGA GAT TGG TCA GTG ACT GGA GTT CCT TGG CAC CCG AGA ATT CCA |
| Multiplexing PCR Reverse Primer PCRId5 | CAA GCA GAA GAC GGC ATA CGA GAT CAC TGT GTG ACT GGA GTT CCT TGG CAC CCG AGA ATT CCA |
| Multiplexing PCR Reverse Primer PCRId6 | CAA GCA GAA GAC GGC ATA CGA GAT ATT GGC GTG ACT GGA GTT CCT TGG CAC CCG AGA ATT CCA |
| Multiplexing PCR Reverse Primer PCRId7 | CAA GCA GAA GAC GGC ATA CGA GAT GAT CTG GTG ACT GGA GTT CCT TGG CAC CCG AGA ATT CCA |
| Multiplexing PCR Reverse Primer PCRId8 | CAA GCA GAA GAC GGC ATA CGA GAT TCA AGT GTG ACT GGA GTT CCT TGG CAC CCG AGA ATT CCA |
| Multiplexing PCR Reverse Primer PCRId9 | CAA GCA GAA GAC GGC ATA CGA GAT CTG ATC GTG ACT GGA GTT CCT TGG CAC CCG AGA ATT CCA |
| Multiplexing PCR Reverse Primer PCRId10 | CAA GCA GAA GAC GGC ATA CGA GAT AAG CTA GTG ACT GGA GTT CCT TGG CAC CCG AGA ATT CCA |
| Multiplexing PCR Reverse Primer PCRId11 | CAA GCA GAA GAC GGC ATA CGA GAT GTA GCC GTG ACT GGA GTT CCT TGG CAC CCG AGA ATT CCA |
| Multiplexing PCR Reverse Primer PCRId12 | CAA GCA GAA GAC GGC ATA CGA GAT TAC AAG GTG ACT GGA GTT CCT TGG CAC CCG AGA ATT CCA |
| Multiplexing PCR Reverse Primer PCRId13 | CAA GCA GAA GAC GGC ATA CGA GAT TTG ACT GTG ACT GGA GTT CCT TGG CAC CCG AGA ATT CCA |
| Multiplexing PCR Reverse Primer PCRId14 | CAA GCA GAA GAC GGC ATA CGA GAT GGA ACT GTG ACT GGA GTT CCT TGG CAC CCG AGA ATT CCA |
| Multiplexing PCR Reverse Primer PCRId15 | CAA GCA GAA GAC GGC ATA CGA GAT TGA CAT GTG ACT GGA GTT CCT TGG CAC CCG AGA ATT CCA |
| Multiplexing PCR Reverse Primer PCRId16 | CAA GCA GAA GAC GGC ATA CGA GAT GGA CGG GTG ACT GGA GTT CCT TGG CAC CCG AGA ATT CCA |
| Multiplexing PCR Reverse Primer PCRId17 | CAA GCA GAA GAC GGC ATA CGA GAT CTC TAC GTG ACT GGA GTT CCT TGG CAC CCG AGA ATT CCA |
| Multiplexing PCR Reverse Primer PCRId18 | CAA GCA GAA GAC GGC ATA CGA GAT GCG GAC GTG ACT GGA GTT CCT TGG CAC CCG AGA ATT CCA |
| Multiplexing PCR Reverse Primer PCRId19 | CAA GCA GAA GAC GGC ATA CGA GAT TTT CAC GTG ACT GGA GTT CCT TGG CAC CCG AGA ATT CCA |
| Multiplexing PCR Reverse Primer PCRId20 | CAA GCA GAA GAC GGC ATA CGA GAT GGC CAC GTG ACT GGA GTT CCT TGG CAC CCG AGA ATT CCA |

| | |
|---|---|
| Multiplexing PCR Reverse Primer PCRId21 | CAA GCA GAA GAC GGC ATA CGA GAT CGA AAC GTG ACT GGA GTT CCT TGG CAC CCG AGA ATT CCA |
| Multiplexing PCR Reverse Primer PCRId22 | CAA GCA GAA GAC GGC ATA CGA GAT CGT ACG GTG ACT GGA GTT CCT TGG CAC CCG AGA ATT CCA |
| Multiplexing PCR Reverse Primer PCRId23 | CAA GCA GAA GAC GGC ATA CGA GAT CCA CTC GTG ACT GGA GTT CCT TGG CAC CCG AGA ATT CCA |
| Multiplexing PCR Reverse Primer PCRId24 | CAA GCA GAA GAC GGC ATA CGA GAT GCT ACC GTG ACT GGA GTT CCT TGG CAC CCG AGA ATT CCA |
| Multiplexing PCR Reverse Primer PCRId25 | CAA GCA GAA GAC GGC ATA CGA GAT ATC AGT GTG ACT GGA GTT CCT TGG CAC CCG AGA ATT CCA |
| Multiplexing PCR Reverse Primer PCRId26 | CAA GCA GAA GAC GGC ATA CGA GAT GCT CAT GTG ACT GGA GTT CCT TGG CAC CCG AGA ATT CCA |
| Multiplexing PCR Reverse Primer PCRId27 | CAA GCA GAA GAC GGC ATA CGA GAT AGG AAT GTG ACT GGA GTT CCT TGG CAC CCG AGA ATT CCA |
| Multiplexing PCR Reverse Primer PCRId28 | CAA GCA GAA GAC GGC ATA CGA GAT CTT TTG GTG ACT GGA GTT CCT TGG CAC CCG AGA ATT CCA |
| Multiplexing PCR Reverse Primer PCRId29 | CAA GCA GAA GAC GGC ATA CGA GAT TAG TTG GTG ACT GGA GTT CCT TGG CAC CCG AGA ATT CCA |
| Multiplexing PCR Reverse Primer PCRId30 | CAA GCA GAA GAC GGC ATA CGA GAT CCG GTG GTG ACT GGA GTT CCT TGG CAC CCG AGA ATT CCA |
| Multiplexing PCR Reverse Primer PCRId31 | CAA GCA GAA GAC GGC ATA CGA GAT ATC GTG GTG ACT GGA GTT CCT TGG CAC CCG AGA ATT CCA |
| Multiplexing PCR Reverse Primer PCRId32 | CAA GCA GAA GAC GGC ATA CGA GAT TGA GTG GTG ACT GGA GTT CCT TGG CAC CCG AGA ATT CCA |
| Multiplexing PCR Reverse Primer PCRId33 | CAA GCA GAA GAC GGC ATA CGA GAT CGC CTG GTG ACT GGA GTT CCT TGG CAC CCG AGA ATT CCA |
| Multiplexing PCR Reverse Primer PCRId34 | CAA GCA GAA GAC GGC ATA CGA GAT GCC ATG GTG ACT GGA GTT CCT TGG CAC CCG AGA ATT CCA |
| Multiplexing PCR Reverse Primer PCRId35 | CAA GCA GAA GAC GGC ATA CGA GAT AAA ATG GTG ACT GGA GTT CCT TGG CAC CCG AGA ATT CCA |
| Multiplexing PCR Reverse Primer PCRId36 | CAA GCA GAA GAC GGC ATA CGA GAT TGT TGG GTG ACT GGA GTT CCT TGG CAC CCG AGA ATT CCA |
| **Filter binding assay for Ago1•let-7** | **Sequence**<br>Seed; m, 2′-*O*-methyl ribose; ps; phosphorothioate |

| | complementary to guide |
|---|---|
| Complete complementary target to *let-7* | pGAUACUAUACAACmCpsmUACUACCUCAACCU |
| Target to *let-7* with seed only pairing (g2g8:t2–t8) | pGAAAAAAAAAAAAmApsmAUCUACCUCUAAAU |
